# Emergent patterns of patchiness reflect decoupling between ocean physics and biology

**DOI:** 10.1101/2024.05.24.595779

**Authors:** Patrick Clifton Gray, Emmanuel Boss, Guillaume Bourdin, Mission Microbiomes AtlantECO, Tara Pacific Consortium, Yoav Lehahn

## Abstract

While a rich history of patchiness research has explored spatial structure in the ocean, there is still no consensus over the controls on biological patchiness and how biogeochemical processes and patchiness relate. The prevailing thought is that physics structures biology, but this has not been tested at the basin scale with consistent *in situ* measurements. Here we show that the patchiness of physics and biology are decoupled at the global scale through analysis of a global dataset of in situ surface optical properties from the S/V *Tara* and using the slope of spatial scale vs variance to quantify patchiness. Based on analysis of ∼650,000 nearly continuous (dx∼150m) measurements from an underway sampling system - representing five years of consistently collected data across the Atlantic, Pacific, and Southern Oceans - we find the patchiness of physical and biological parameters are uncorrelated. We show that variance slope is an emergent property with unique patterns in biogeochemical properties that are distinct from physical tracers, yet connected to other biological tracers. These results provide context for decades of discrepancy between *in situ* studies, could support new tests of biogeochemical model parameterizations, and open the way for new insight into processes regulating the observed patterns.

## Introduction

Most phytoplankton species near the ocean’s surface have a doubling time on the order of a day^1^. This rapid doubling time means phytoplankton are extremely responsive to nutrient fluxes and external forcing^2^. Coupled with the dynamic physical nature of the marine environment and grazing pressures with resulting loss rates similar to growth rates for phytoplankton^3,4^, this results in considerable spatial heterogeneity, or patchiness, at all scales^5–8^. This spatial heterogeneity supports increased diversity by creating different environmental conditions within close proximity, ecosystem stability through diversity and maintenance of seed populations, and efficient transfer of carbon up trophic levels^9^. The link between patchiness and the efficient transfer of carbon in the marine ecosystem is largely due to dense aggregations that make it energy efficient for zooplankton to graze, and patches of zooplankton make it feasible for planktivorous fish to meet their own energy requirements^10,11^. This spatial variability of phytoplankton and zooplankton, and the generally increasing patchiness as one moves up the trophic chain, has important implications for food availability and productivity of the entire ocean food web^12,13^.

The spatial patterns of the plankton ecosystem and their drivers have long been investigated. An early outline by Hutchinson discusses nutrient and temperature gradients, stochastic events such as storms, intra-species signaling, competition, and predation among processes that likely control these patterns^14^. An updated synthesis by Levin described pattern and scale as *the* central problem in ecology, arguing for cross-scale investigations to understand the mechanisms and consequences of marine patchiness^5^. The scale-dependent spatial heterogeneity exhibited by both biology and physics has been a central focus of studies into pattern and scale in the ocean^6^, with much of this work attempting to divide spatial scales into domains of scale-independence and then investigating the dominant processes influencing biology in those domains^15,16^. Power spectral analysis, a Fourier decomposition where variance is partitioned into the contribution of bands of specific frequencies or wavenumbers (see ^17^ for an early review on the implementation of the technique in ecology), is often used to describe phytoplankton patchiness and quantify this scale-dependence. The scaling behavior of a geophysical variable is typically represented as a log-log plot of variance against length scale. If that relationship is approximately linear (in log space) this represents a power law relation between variance and spatial scale where the exponent of that power law relationship is the spectral slope^18^. A flatter spectral slope indicates less of a decrease of variance with scale, i.e. more variability at smaller scales relative to larger scales, when compared to a steeper slope. This parameter has often been used synonymously with patchiness, but can also be thought of as quantifying the cascade of variance across scales. While there are a range of issues with this decomposition, such as discarding of phase information^9,19^, it is commonly used as an indicator of the distribution of spatial variance.

There is over a century of inquiry into the spatial heterogeneity of ocean physics. Turbulence in the ocean generally cascades down spatial scales. Turbulent mesoscale eddies spin off of large-scale circulation features. Energy then transfers from these major eddies to smaller turbulent whorls and finally to viscous dissipation into heat^20,21^. This turbulent cascade generally has a spectral slope of -5/3 in the inertial subrange (a spatial range smaller than the major energy containing eddies, but larger than viscous eddies). While Kolomogrov initially proposed this cascade of variance for fully developed 3D turbulence in the ocean’s inertial subrange, O(10-0.001) meters depending on flow characteristics^22^, the -5/3 spectral slope has been shown to hold generally true for the inertial subrange of 2D ocean turbulence O(100-1) km as well^23^. Temperature generally follows the same cascade as turbulent kinetic energy. It has a source of variance at large scales, a dissipation of variance at small scales, and it is typically being mixed by similar turbulent processes^24^.

The patchiness of phytoplankton was initially attributed simply to turbulent stirring by physical processes^25^, with a general conclusion of consistent scaling behavior between physical and biological features^16^. This conclusion was drawn largely from its spectral slope that appeared similar to the -5/3 of 2D turbulence^25,26^. Modeling indicated that the spectral slope of phytoplankton was also influenced by the phytoplankton reproductive rate, diagnosed by a scaling break at larger spatial scales where the spectral slope became flatter than that of a passive tracer^27,28^. This led to a substantial amount of work, with many conflicting results, investigating the time and length scales where the contribution of biologically generated spatial variability (growth rates, grazing, etc) to the total spatial variability is significant vs where where physics dominates^6,7,9,22,29^.

Abrahams^7^ suggested and others have corroborated in modeling^30,31^ that the spatial patterns in plankton are a product of the timescales of their response to physically driven changes. For example, the equilibration of temperature to a new heat flux is considerably slower than the response of phytoplankton to a nutrient flux. A small burst of cold nutrient-rich water thus leads to increased spatial heterogeneity in chlorophyll-a (chl-a) vs temperature, but as nutrients are exhausted and the heat is mixed over time chl-a and temperature would return to covarying^29^. Modeling work shows a steeper spectral slope in temperature compared to chl-a, attributing some of the slope variability to low frequency physical variations and the correlation of physical and biological variables to “the dominant role of physics in the spatial variability of plankton distribution”^30^.

Despite these theoretical advances, actual observations remain inconsistent. Some work concludes that at <1km scales biological processes are important for patch dynamics^10,32^, while other work asserts that at this scale spatial distributions of phytoplankton are dominated by turbulent diffusion. For example, this latter conclusion is advanced in work that found a strong correlation between chl-a and temperature patchiness below 5km yet not so above 5 km to 80km^17^. Instances spring up where phytoplankton are patchier than zooplankton^33^, directly conflicting generally accepted findings that the patchiness of phytoplankton is between that of zooplankton and physical factors^7,25,34^. Work assessing the spectral slope of chl-a in the same region for multiple years found completely contrasting measurements from month to month^22,35^. After decades of work, it is not clear at which spatiotemporal scales biological processes are important, or even which processes are dominant in controlling patchiness in the ocean.

While biophysical ocean patchiness has been relatively well documented on local scales, its global characteristics have not yet been analyzed. Therefore, there is little data to test new theories, constrain impacts on productivity and diversity, and simply describe the patchiness itself. Importantly, this limits our ability to parameterize global biogeochemical models that are computationally limited from explicitly modeling these scales in our global models of the carbon cycle.

Here we address this gap, and explore the global characteristics of biophysical patchiness using a unique dataset of *in situ* surface plankton-related optical properties from the S/V *Tara* program. All data comes from the Tara Pacific^36^ and Tara Microbiome missions which represent five years of observations spanning the Atlantic, Pacific, and Southern Oceans. This data was collected and processed consistently. After conservative quality control this dataset amounts to N=661,552 (*d*t=1 minute, *d*x∼150m) measurements used in this analysis (Figure 1). As we aim to discern between patterns of physical and biological patchiness, the dataset variables are divided into two corresponding groups, with the physical domain being represented by temperature, salinity and density, and the biological domain by chl-a, particulate attenuation of light at 443nm (c_p_(443)), and γ an optical proxy for mean particle size^37^. The latter two, because they may include non-living particles, are referred to as biogeochemical properties.

**Figure 1:**
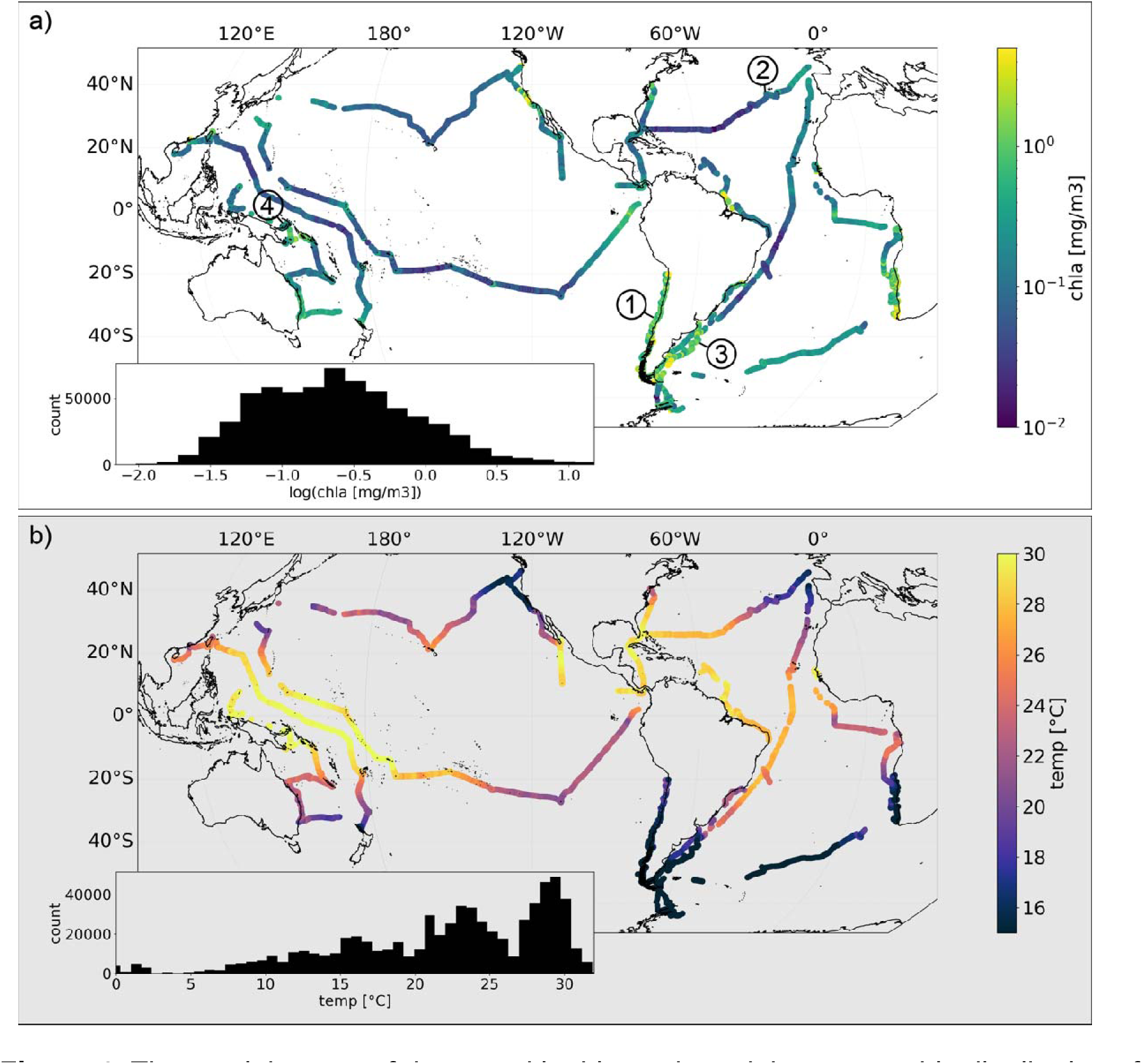
The spatial extent of data used in this study and the geographic distribution of chlorophyll-a (a) and sea surface temperature (b). The data includes 661,552 minute binned measurements. Insets show the histograms of both variables. The four numbers in panel (a) correspond to the example data shown in Figure 2.

## Results and discussion

While the Fourier-based spectral slope is an extremely useful and commonly used tool for characterization of patchiness, it is not suitable for analyzing unevenly sampled datasets. To overcome this limitation here we quantify patchiness via the dependence of the variance, V, on length scale, L following ^31^. As found in previous work, V varies mostly linearly with L in log space (Figure 2), indicating a power law relationship of the form: (1) V = L^I’^, where r describes the slope (exponent) of the relationship in log space. In this work we focus on r, the “variance slope”. As with a power spectral density slope, a transect with a lower r would have relatively more variance retained at smaller length scales relative to larger scales compared to a transect with higher r, i.e. a lower r is patchier. This metric can be thought of qualitatively as a ratio of large scale to small scale variance (Figure S1), with r increasing as the large scale variance dominates the small. This parameter is similar and strongly correlated with the conventional Fourier-based spectral slope exponent (Figure S2 and S3), but our approach (from ^31^) has the advantage of being applicable to data with gaps, as long as they are smaller than 20% of a transect.

**Figure 2.**
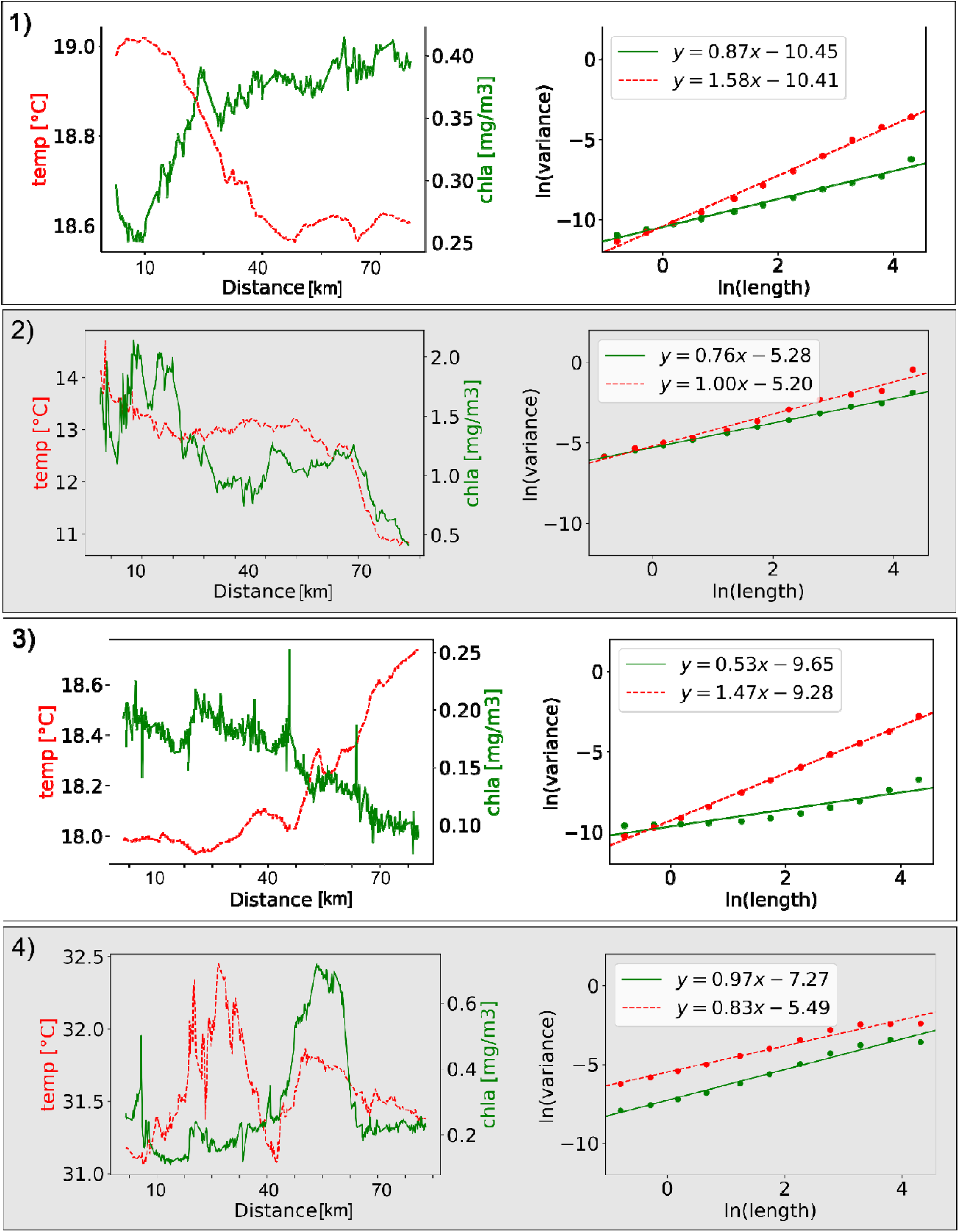
Examples of the chlorophyll-a (green) and temperature (red) absolute values along a leg (left panel in each pair) and the scale-dependent variance used to calculate the variance slope (r) derived from these legs (right panel in each pair). The numbers of each panel (1-4) correspond to the locations in Figure 1a.

Specifically in this work, variance was calculated on subsets of 500 samples over a set of 11 log-distributed windows from 3 to 500 samples, spanning approximately 0.5km to 75km. Both variance and window size were log-transformed, and we ran a least-squares fit to find the slope of this line, yielding the r of V = L^I’^. In our analysis of global patterns of oceanic patchiness, we distinguish between r of the biological and biogeochemical variables chl-a, c_p_(443) and γ ( r_chl-a_, r_cp_ and rγ, respectively), and of the physical variables temperature, salinity and density ( r_temperature_, r_salinity_ and r_density_, respectively).

Comparison between the geographic distributions of r_chla_ and r_temperature_ reveals distinct differences between global patterns of physical and biological patchiness (Figure 3). Notably, the geographic distributions of r_temperature_ are more random, while r_chla_ is consistently lower in oligotrophic regions, particularly the subtropical gyres, and higher in physically energetic regions such as western boundary currents, eastern boundary upwelling zones, and equatorial upwelling zones. In agreement with most previous observations, r_chla_ is, on average, lower than r_temperature_, indicating chl-a is patchier than temperature. r_temperature_ is fairly normally distributed around 1, while r_chla_ is slightly right-skewed with a mean around 0.5 (see inserts in Figure 3). Based on the observed relationship between and the power spectral density slope (Figure S3) the classic -5/3 slope found in previous studies for the inertial subrange would correspond to a value of 0.84 which is near, but slightly lower than the mean of our global _temperature_ calculations, 0.97.

**Figure 3.**
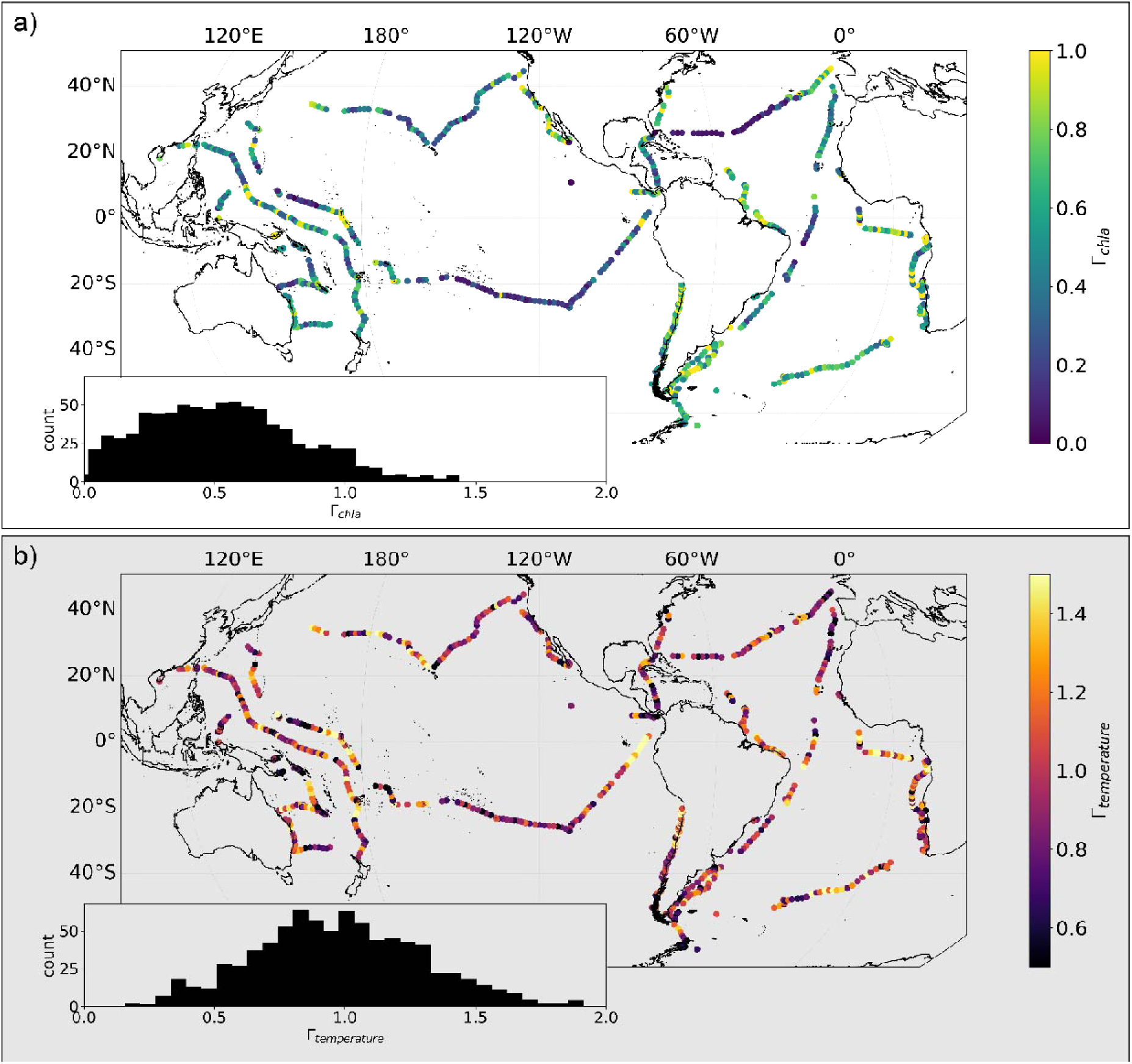
Geographic distributions of _chla_ (a) and _temperature_ (b). Insets show the histograms for these two variables.

Interestingly, running the same analysis on satellite-based chl-a and temperature reveals contradicting results. Although satellite imagery has been successfully used to track individual chl-a patches^38,39^, and to quantify patchiness at local scales^25^, in the satellite-based analysis we find r_temperature_ is lower than r_chla_ (i.e. the opposite of our *in situ* results), and the geographic patterns for both chl-a and temperature are different (Figures S4 and S5, respectively). We attribute this discrepancy primarily to issues of measurement sensitivity and atmospheric correction of ocean color remote sensing (from which chl-a is derived) which is applied as a pixel level correction, artificially injecting variance at small scales. This effect is likely to have the strongest impact in oligotrophic regions, where chlorophyll-a levels are extremely low, resulting in minimal absorption and reduced particulate scattering, and thus a low signal to noise ratio in the satellite retrieval. This could cause issues in both spatial and temporal analyses^40^. This corroborates results from previous work that used a semivariogram approach to analyze variance in global satellite chl-a patterns from SeaWIFS^41^ which showed that unresolved variance was negatively correlated with mean chl-a and the authors suggested the high fraction of unresolved variance in the subtropical gyres might stem from atmospheric correction, though they did not differentiate between instrument noise and submesoscale variability. A more recent, but similar study compared the unresolved variance between MODIS-Aqua and SeaWIFS and found it was reduced, but not eliminated with MODIS^42^.

The global patterns characterizing our observations point to decoupling between the governing processes underlying the formation of physical and biological ocean patchiness. This decoupling is emphasized when plotting the correlation between the different types of patchiness (Figure 4), which we show both for all individual legs and averaged by Longhurst provinces^43^ (Figure 5). The r of biological variables are correlated globally, even between chl-a and y, the mean particle size (Figure 4a,b). Similarly, r of physical variables are well correlated with each other (Figure 4c,d). Yet, there is little to no correlation between the r of biological variables and physical variables (Figure 4e,f). This suggests that ecosystem patchiness is an emergent property due to biological processes rather than driven by physical forcings.

**Figure 4.**
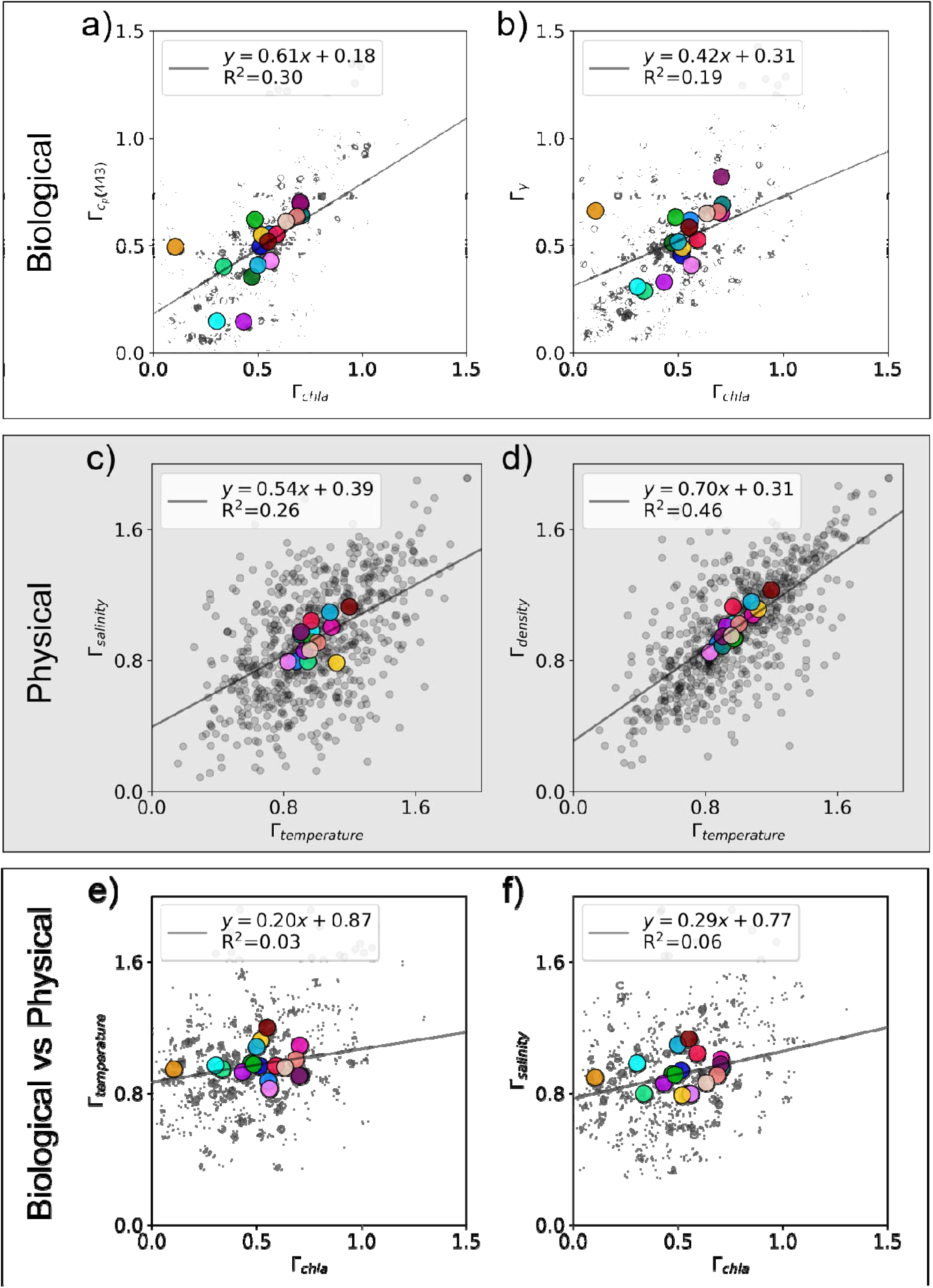
Variance slope () comparisons between different variables. _chla_ correlates with the biogeochemical parameters such as attenuation (a) and (mean particle size) (b), and of different physical variables are correlated (d, c), yet biogeochemical vs physical variables don’t have a notable correlation (e, f). Larger colored markers correspond with the Longhurst provinces from Figure 5. N.b. correlations are shown for all data, not the Longhurst province means, and all plots have a p-value < 0.001.

**Figure 5.**
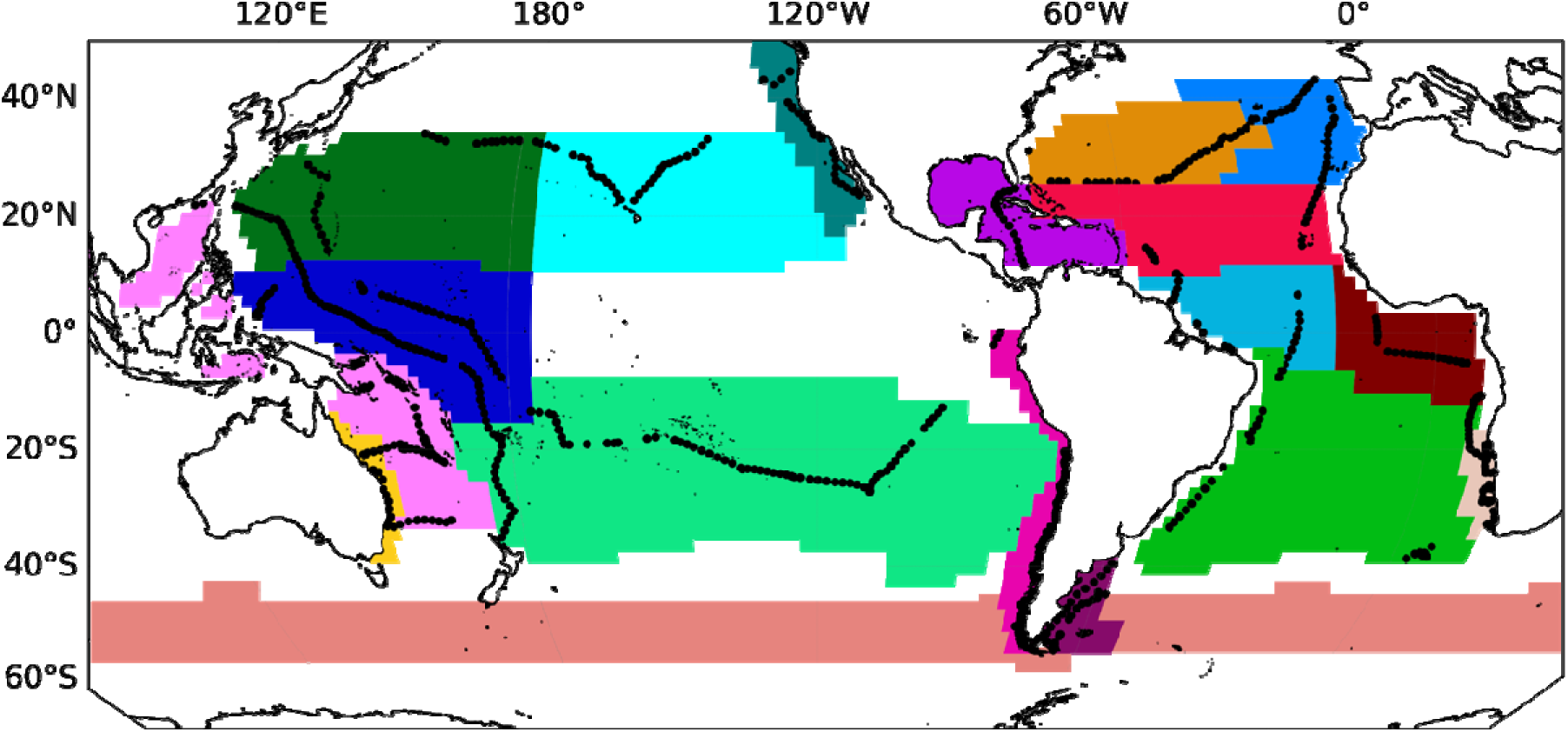
Longhurst provinces that contain N=> 25 legs shown geographically along with the Tara legs within each.

If we decrease the top-end length scale that is used for the calculation of r, for example from 75km to 5km, r_chla_ and r_temperature_ become only marginally more correlated (R^2^=0.04 for 0.5km to 5km vs R^2^=0.03 for 0.5km to 75km). This is in contrast with older work which showed that chl-a and temperature patchiness were strongly correlated below 5km yet not so above 5 km to 80km^17^.

Similarly to the differences between geographic distributions patterns (Figures 1 and 3) the correlations between r of different physical variables differ substantially from the correlations between absolute values (Figure S6). We show chl-a and gamma have minimal correlation in absolute value yet a correlation in r, chl-a and temperature have a strong correlation in absolute value yet minimal in r, and temperature and salinity have minimal correlation in absolute value yet a strong correlation in r.

Another fundamental difference between global patterns of physical and biological ocean patchiness emerges when looking at the data averaged by biogeochemical provinces and plotting the relationship between and the absolute value of each variable (Figure 6). The biological variables that are concentration dependent (i.e. chl-a and cp(443)) have a positive relationship between their and their absolute value, such that high biomass provinces are less patchy (i.e. associated with higher) than low biomass provinces, (Figure 6a-c). In contrast, there is no relationship between and the absolute value of the physical variables (Figure 6d-e). In other words, biomass and biological patchiness are linked (inversely) while heat and salt content does not correlate with their patchiness.

**Figure 6.**
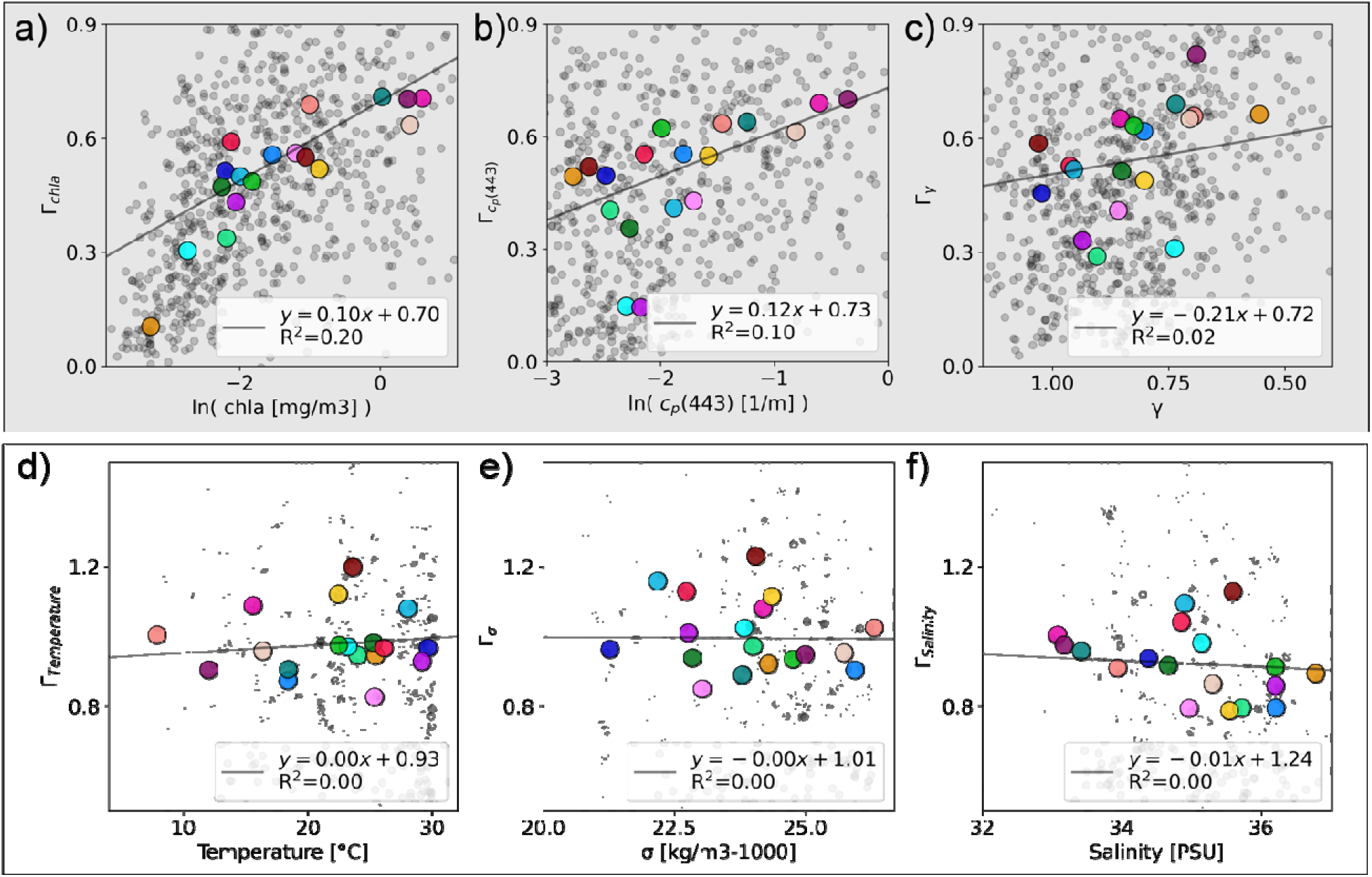
The top (a-c) and bottom (d-f) rows show the absolute value vs variance slope () for a range of biological and physical variables, respectively. The average of each province is shown in the larger markers and the colors correspond to the provinces on the map. N.b. correlations are shown for all data, not the Longhurst province means, and all plots have a p-value < 0.001.

The linkage between biological patchiness and biological productivity is emphasized when taking into account nutrient availability averaged by Longhurst Province (Figure 7). When mapped onto the absolute value of chl-a and _chla_, a range of nutrients have a clear increasing relationship with both (Figure 7), with the highest nutrient values also having high chl-a and _chla_.

**Figure 7.**
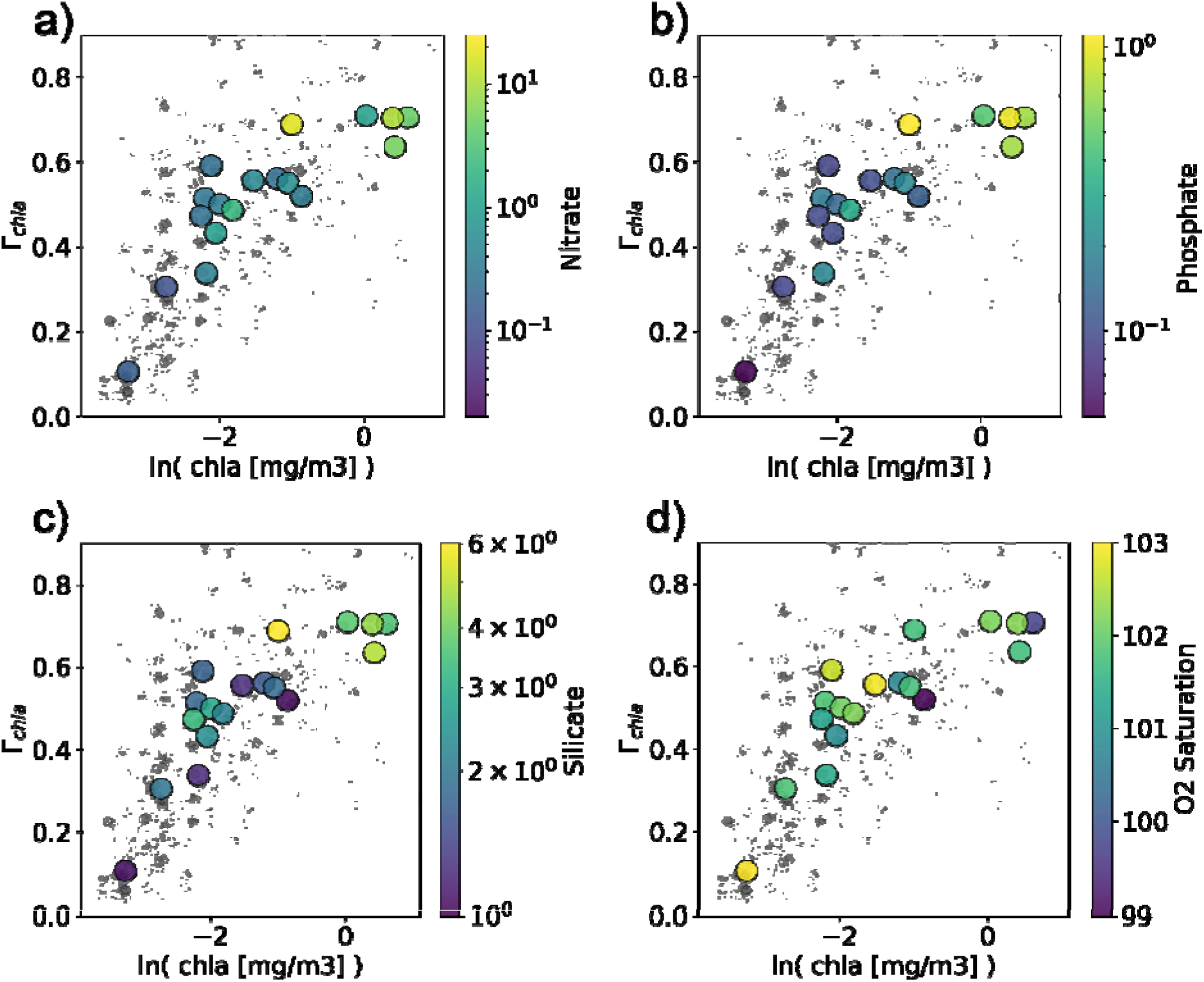
Chlorophyll-a (chl-a) concentration vs _chla_ colored by average nutrient concentrations in each Longhurst province. Nutrient data is from the World Ocean Atlas 2018.

The nature of the small scale variability in biological signals, and the way they differ from the physical signals, is emphasized when zooming on segments of the transects (Figure 8). Careful examination reveals that while the high frequency changes in our underway observations of biological variables may appear to be associated with instrumental noise, the individual measurements are robust, and the changes reflect small scale (<1km) biological variability patterns (Figure 8). This robustness can be seen both in the covariation between different sensors (Figure 8a) and the raw absorption and attenuation spectra (Figure 8b,c). This small scale variability lowers and can make the variance spectra nearly flat in regions without low frequency variation, such as the Sargasso Sea and Subtropical South Pacific.

**Figure 8.**
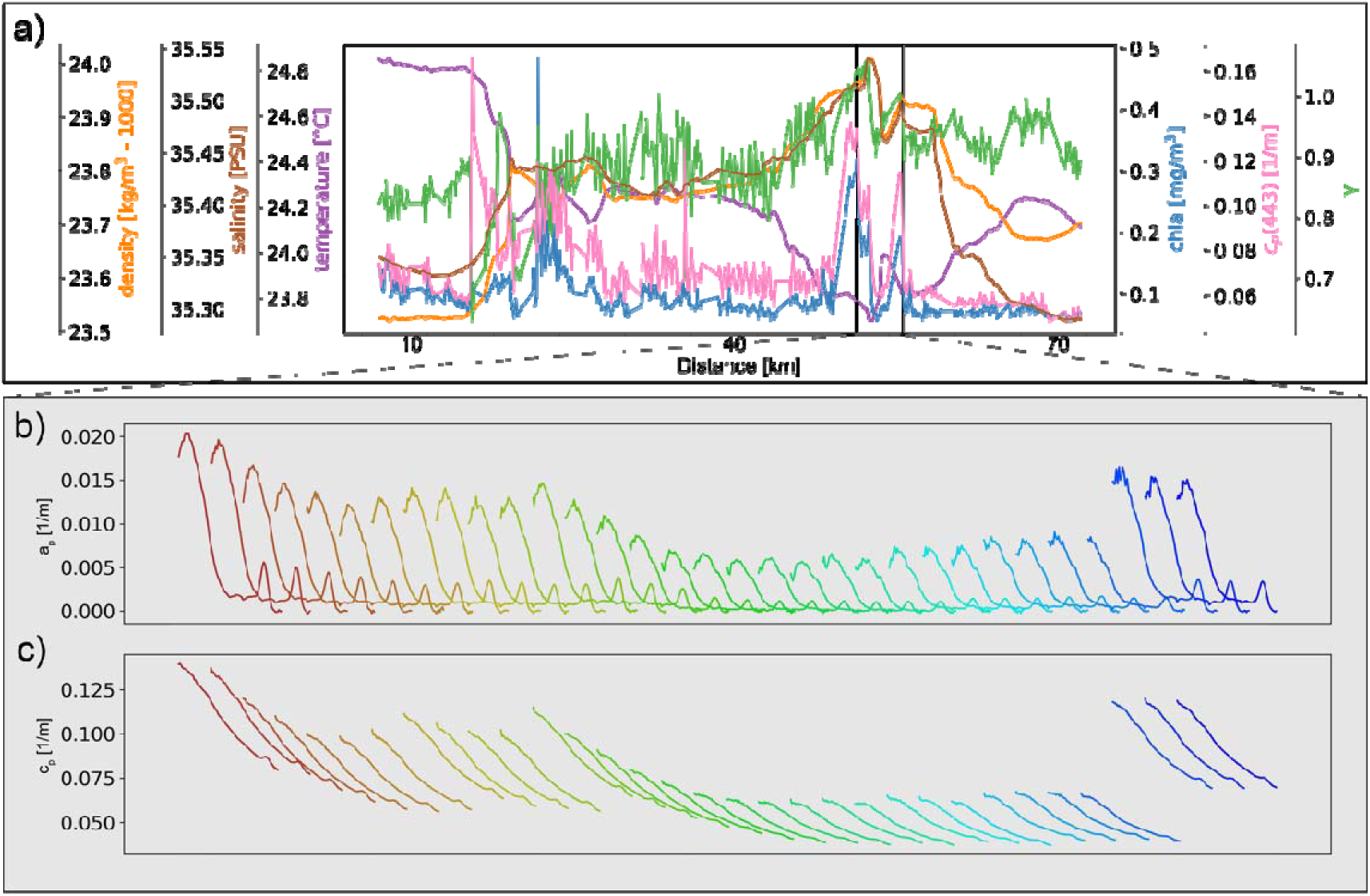
Example of measured variables along a single leg (a), showing data “spikes” that may initially be considered instrumental noise, but when inspecting the individual spectra (b and c), used to derive the various proxies, they are consistent with expectation and across the absorption and attenuation sensors. The two bottom rows show a 30 minute subset of particulate absorption spectra (b) and particulate attenuation spectra (c) representing the period between the vertical black lines in panel a. Each spectra is the average of one minute of sampling, is colored by time (from brown to blue), and spans wavelengths from 410nm to 750nm. This transect is from the Coral Sea.

Our work does not tie the variance slope to any particular process. Possibly r represents the integrated impact of physical and biogeochemical characteristics within a region. The lower r_chla_ in oligotrophic regions may be attributed to a sustained injection of variance from ecosystem interactions (e.g. growth, grazing). Recent work investigating zooplankton patchiness found coherent small scales patches across many taxa dominated by length scales of 10-30 m^40^. Even highly idealized models have been shown to generate patchiness via grazing^44^ which could help explain aspects of our results. While previous work using this same metric suggested that the response time of phytoplankton (i.e. growth rate) is faster than the response time of temperature (i.e. equilibration time) as the reason chl-a is patchier^31^, this doesn’t agree with our results, where the regions with higher assumed growth rates are less patchy (e.g. eastern upwelling zones). This may suggest that certain elements of the ecosystem are responding faster in these highly patchy oligotrophic regions, possibly grazers or growth following fine-scale vertical nutrient fluxes. Stratification may play a role with a more stratified water column leading to more intermittent linkages between the surface and the nutrient-rich deep ocean, resulting in turn in a patchier expression in the highly stratified subtropical regions. While the drivers are not clear from our work, Mahadevan & Campbell^31^ suggest that an increasingly patchy distribution requires finer scale sampling or grid spacing, and if these processes are important, this indicates fine-scale grid spacing is in fact still important across broad swaths of the ocean generally considered “homogenous”.

While the focus of this work is on scaling relationships, the fractal-like patchiness of phytoplankton is not infinite. At a fundamental level, patches are formed by individual phytoplankton living and dying^45^. Currently it is unclear how to connect the scales being investigated here with the individual organisms at the heart of the matter; Franks^22^ notes that it is not likely we can diagnose the dynamics underlying the observations from spectral slopes alone. Towards mechanistic understanding of the processes underlying the spatial patterns we expect Lagrangian approaches will be key, particularly to parse out spatial and temporal dynamics. Future work should examine joint spatial patterning across trophic levels, combining the methods here, rapid methods for a wide size range of plankton^46^, and methods for quantifying zooplankton patchiness^10^. Additionally, it may be that improved atmosphere correction of satellite products could enable a new source of information via spatial analyses across the global ocean. Higher signal to noise regions, such as coastal waters or higher latitudes during the spring bloom, may be more amenable to satellite analyses of spatial patterns under existing atmospheric correction schemes.

## Summary and conclusion

We identified two fundamental differences between global patterns of physical and biological patchiness: globally biological patchiness at the scales analyzed here (∼0.5km to 75km) is itself spatially organized while physical patchiness is more random, and patchiness of concentration-dependent biological variables is correlated with the absolute values of these variables, while no such correlation is found for physical variables. The statistical distributions of r_chla_ and r_temperature_ agree with previous work, where r_chla_ has a generally lower value, i.e. a patchier spatial pattern, compared to r_temperature_. We find global patterns of biological patchiness to coincide with biogeochemical provinces and with nutrients levels. While our analysis shows the patchiness of biological variables are intercorrelated, and patchiness of physical variables are intercorrelated, contrary to common view, we find no interdependence between physical and biological patchiness. Our results point to decoupling between the governing processes underlying the formation of physical and biological ocean patchiness. This suggests that biological processes regulate the spatial patterning of the oceanic ecosystem independently of the physical processes or, rather than true independence, that they modulate the initial physical-driven spatial pattern baseline sufficiently to manifest as a decoupling. We emphasize here that the absolute value of variance itself is correlated at all scales between physical and biological variables (Figure S7). Moreover, while it is well known that there is a relationship between temperature and chl-a, largely attributed to nutrient fluxes^47^, our findings show this does not extend to variance slope. Our results contradict results from model studies (e.g. ^30^) and from satellite analyses (e.g. ^48^), including our own (Figure S4 and S5). This discrepancy holds even when comparing to other work on the same range of scales as those examined here, which had concluded that biological patchiness is controlled by mesoscale mixing^25^. This suggests that these two important tools for understanding ocean ecosystems (models and satellite observations), fail to capture fundamental characteristics of the marine ecosystem. Our results may also help explain the disagreement shown across decades of patchiness research including many *in situ* studies. It may be that there is no generalizable patchiness relationship between physics and biology in the ocean and no consistent spatiotemporal scale where biological or physical processes dominate the spatial patterns of phytoplankton.

Accordingly, basing general theories on the patchiness of a given tracer type from a specific site and over short time scales could be misguided. Instead, the ocean appears to consist of myriad physical and biogeochemical processes manifesting as a diverse set of spatial patterns. Thus, while no consistent relationships emerge these observable spatial patterns may provide a window into the processes creating them.

These results support the use of low dimensional spatial encodings such as r to study these upper-ocean processes. We speculate these spatial metrics may be sensitive to biogeochemical parameters not represented by the absolute value of chl-a. Importantly, we cannot resolve marine ecosystems at the mesoscale and below in basin-scale biogeochemical models, thus we must increase our ability to observe, invert, and understand what our models are missing^49^. Spatial patchiness patterns may help drive our models to reality and serve as an underutilized source of insight into the underlying processes generating the observed patchiness.

## Methods

All *in situ* measurements are from the S/V *Tara* underway sampling system. This pulls seawater from 2m depth. Temperature and salinity data is from a Seabird Scientific thermosalinograph (SBE 38 and SBE 45) and all optical data is derived from particulate absorption and attenuation measured with a Seabird Scientific ACs.

We quantify patchiness using variance slope, r, following ^31^ which is similar to spectral analysis, but able to ingest vectors (i.e. transects) with small gaps within them. r is calculated on groups of 500 samples using a set of 11 log-distributed windows from 0.5km to 75km. Variance and window size were log-transformed, and we ran a least-squares fit to find the slope of this line, yielding the r of V = L^I’^. It is worth noting a few characteristics of r to gain some intuition into the metric. The variance slope of a randomly distributed variable is centered around 0. The calculation is not sensitive to whether a variable is log-normally distributed (such as chl-a). For example, r_chla_ is identical whether raw chl-a values are input or log-transformed chl-a values. The S/V *Tara* is a sailing vessel and thus does not always travel in a straight line or with the same speed so here we have used time (minute bins of data) instead of distance to calculate the window, but filtered out legs that have a maximum distance traveled under 30km or over 150km. We have required at least 80% of the transect to contain data and filtered out transects where the maximum time elapsed is greater than 16 hours.

Hyperspectral absorption and attenuation (400 to 735 nm at ∼4 nm spectral resolution, AC-s, Seabird Sci.) were measured continuously. A 0.2 µm filter cartridge was connected to the system and we automatically redirected the flow to measure the properties of filtered seawater for 10 minutes every hour. Total (“normal”) seawater was flowing the rest of the time. Absorption and attenuation measurements recorded during the filtered periods were interpolated across the transect and subtracted from the total seawater measurements to obtain an estimate of the particulate absorption (a_p_), attenuation (c_p_) spectra. During the Tara Microbiome transect, the filtered periods were interpolated using the variation in CDOM fluorescence (fCDOM) also measured continuously with a SeaPoint ultraviolet fluorometer (SUVF). fCDOM data was recorded with a Seabird Scientific WSCD CDOM fluorometer during the Tara Pacific transect.

The signal to noise ratio of the WSCD is lower than the SUVF’s which introduced noise in the fCDOM interpolation and the product derivation. Therefore, the filtered periods were interpolated linearly across the Tara Pacific transect. This approach permits retrieval of particulate optical properties independently from the instrument drift and biofouling^50^. For each minute of total seawater measurements (sampled at 4Hz) the signal between the 2.5th and 97.5th percentiles were averaged, and their standard deviation was used to quantify uncertainty. Dropping the 2.5th to 97.5th percentiles filters out noisy spikes from bubbles which can be a major problem in optical measurements. All spectra were manually checked and quality controlled for obviously bad measurements (e.g. bubbles, bad filtered seawater measurements). These inherent optical properties were used as proxies for a range of particulate properties, primarily chl-a line height, a chl-a estimate derived from the absorption peak at 676 nm^51,52^, and y, a proxy for mean particle size^37^.

Underway data was collected using Inlinino^53^ an open-source logging and visualization program, and processed using InlineAnalysis (https://github.com/OceanOptics/InLineAnalysis) following best practices^54^.

Longhurst provinces ^43^ (downloaded from https://www.marineregions.org/sources.php#longhurst) were used as approximate delineations of the global ocean into biogeochemical regions. Nutrient data was from the World Ocean Atlas 2018 (downloaded from https://www.ncei.noaa.gov/data/oceans/woa/WOA18).

## Data Availability

All data is available in raw form at NASA’s SeaBASS archive via the search keyword Tara_Microbiome. All data prepared and formatted for this study is available on GitHub https://github.com/patrickcgray/spatial_patchiness_tara, easily ingestable as a geofeather at multiple stages of processing.

## Code Availability

All code necessary to generate the figures is on GitHub https://github.com/patrickcgray/spatial_patchiness_tara. To obtain a Docker image for running this in an identical environment to the one used in this study run “*docker pull pangeo/pangeo-notebook:2024.04.05*” from the command line with Docker Desktop running. A series of Jupyter Notebooks are provided in the above Github repo for an exact reproduction of all work in this study.

## Acknowledgements

We acknowledge support from the Zuckerman STEM Leadership Program to PCG. NSF EarthCube program award #2026932 supported the Pangeo Cloud platform where all analysis was conducted. The optical inline dataset was collected and analyzed with support from NASA Ocean Biology and Biogeochemistry program under grants NNX13AE58G, NNX15AC08G and 80NSSC21K0783, and 80NSSC20K1641 to the University of Maine. We wish to thank the Tara Ocean Foundation, the SV Tara crew and all those who participate in Mission Microbiomes AtlantECO and adopt its Data Sharing & Publication Best Practices (https://zenodo.org/communities/mission-microbiomes-atlanteco/). This publication has received funding from the European Union’s Horizon 2020 research and innovation programme under grant agreement No 862923 (project AtlantECO). This output reflects only the author’s view and the European Union cannot be held responsible for any use that may be made of the information contained therein. We are keen to thank the commitment of the following institutions for their financial and scientific support that made Mission Microbiomes AtlantECO possible: Stazione Zoologica Anton Dohrn, European Bioinformatics Institute (EMBL-EBI), Centre national de la recherche scientifique (CNRS), Centre National de Séquençage (CNS, Genoscope), agnès b., BIC, Capgemini Engineering, Fondation Groupe EDF, Compagnie Nationale du Rhône, L’Oréal, Biotherm, Région Bretagne, Lorient Agglomération, Billerudkorsnas, Havas Paris, Fondation Rothschild, Office Français de la Biodiversité, AmerisourceBergen, Philgood Foundation, UNESCO-IOC, Etienne Bourgois. Special thanks to the Tara Ocean Foundation, the S/V Tara crew and the Tara Pacific Expedition Participants (https://doi.org/10.5281/zenodo.3777760). We are keen to thank the commitment of the following institutions for their financial and scientific support that made this unique Tara Pacific Expedition possible: CNRS, PSL, CSM, EPHE, Genoscope, CEA, Inserm, Université Côte d’Azur, ANR, agnès b., UNESCO-IOC, the Veolia Foundation, the Prince Albert II de Monaco Foundation, Région Bretagne, Billerudkorsnas, AmerisourceBergen Company, Lorient Agglomération, Oceans by Disney, L’Oréal, Biotherm, France Collectivités, Fonds Français pour l’Environnement Mondial (FFEM), Etienne Bourgois, and the Tara Ocean Foundation teams. Tara Pacific would not exist without the continuous support of the participating institutes. The authors also particularly thank Serge Planes, Denis Allemand, and the Tara Pacific consortium.

The members of the Mission Microbiomes AtlantECO are as follows: Bourdais A. (1), Bowler C. (2), Moulin C. (1), de Vargas C. (3), Iudicone D. (4), Couet D. (3), Catafort E. (5), Boss E. (6), Petit E. (7), Mayeux E. (7), Lombard F. (3), Schramm J. (1), Guidi L. (3), Moll M. (8), Wincker P. (7), Laxenaire R. (9), Troublé R. (1), Sanchez S. (7), Pesant S. (10), Linkowski T. (1) with (1) Tara Ocean Foundation, France; (2) Centre national de la recherche scientifique, France; (3) Sorbonne Université, France; (4) Stazione Zoologica Anton Dohrn, Italy; (5) World Courier, France; (6) School of Marine Sciences, University of Maine, USA; (7) Centre de l’énergie atomique, France; (8) EMS Sistemas, Spain; (9) Ecole Normale Supérieure, France; (10) European Molecular Biology Laboratory, Germany The members of the Tara Pacific Consortium are as follows: Planes S. (1), Allemand D. (2), Agostini S. (15), Armstrong E. (20), Audrain S. (12), Aury J.-M. (20), Banaig B. (1), Barbe V. (20), Belser C. (20), Beraud E. (2), Boissin E. (1), Bonnival E. (22), Boss E. (13), Bourdin G. (13), Bourgois E. (12), Bowler C. (25), Carradec Q. (20), Cassar N. (27, 28), Cohen N. R. (30), Conan P. (10), Cronin D. R. (19), da Silva O. (17), de Vargas C. (22), Djerbi N. (3), Dolan J. R. (17), Dominguez Herta G. (19), Douville E. (14), Du J. (19), Filée J. (32), Flores J. M. (8), Forcioli D. (3), Friedrich R. (26), Furla P. (3), Galand P. E. (11), Ghiglione J.-F. (10), Gilson E. (3), Gorsky G. (17), Guinther M. (15), Haëntjens N. (13), Henry N. (22), Hertau M. (12), Hochart C. (11), Hume B. C. C. (4), Iwankow G. (1), John S. G. (29), Karp-Boss L. (13), Kelly R. L. (29), Kitano Y. (16), Klinges G. (21), Koren I. (8), Labadie K. (20), Lancelot J. (12), Lang-Yona N. (9), Lê-Hoang J. (20), Lemee R. (17), Lin Y. (27), Lombard F. (17), Marie D. (22), McMind R. (3), Miguel-Gordo M. (24), Trainic M. (8), Monmarche D. (12), Moulin C. (12), Mucherie Y. (12), Noel B. (20), Ottaviani A. (3), Paoli L. (7), Pedrotti M.-L. (17), Pesant S. (23), Pogoreutz C. (4), Poulain J. (20), Pujo-Pay M. (10), Reverdin G. (31), Reynaud S. (2), Romac S. (22), Röthig T. (5), Rottinger E. (3), Rouan A. (3), Ruscheweyh H.-J. (7), Salazar G. (7), Sullivan M. B. (18), Sunagawa S. (7), Thomas O. P. (24), Troublé R. (12), Vardi A. (9), Vega-Thunder R. (21), Voolstra C. R. (4), Wincker P. (20), Zahed A. (19), Zamoum T. (3), Ziegler M. (6), and Zoccola D. (2) on behalf of the consortium Tara Pacific. With (1) PSL Research University: EPHE-UPVD-CNRS, USR 3278 CRIOBE, Université de Perpignan, France; (2) Centre Scientifique de Monaco, Principality of Monaco; (3) Université Côte d’Azur, CNRS, Inserm–IRCAN; (4) Department of Biology, University of Konstanz, Konstanz, Germany; (5) Aquatic Research Facility, Environmental Sustainability Research Centre, University of Derby, Derby, United Kingdom; (6) Department of Animal Ecology & Systematics, Justus Liebig University, Giessen, Germany; (7) Department of Biology, Institute of Microbiology and Swiss Institute of Bioinformatics, ETH Zürich, Zürich, Switzerland; (8) Weizmann Institute of Science, Dept. Earth and Planetary Science, Israel; (9) Weizmann Institute of Science, Dept. Plant and Environmental Science, Israel; (10) Sorbonne Université, CNRS, LOMIC, Observatoire Océanologique de Banyuls; (11) Sorbonne Université, CNRS, LECOB, Observatoire Océanologique de Banyuls; (12) Tara Ocean Foundation, Paris, France; (13) School of Marine Sciences, University of Maine, United States of America; (14) Laboratoire des Sciences du Climat et de l’Environnement, LSCE/IPSL, CEA-CNRS-UVSQ, Université Paris-Saclay, France; (15) Shimoda Marine Research Center, University of Tsukuba, Shizuoka, Japan; (16) National Institute of Environmental Science, Japan; (17) Sorbonne Université, Institut de la Mer de Villefranche sur mer, Laboratoire d’Océanographie de Villefranche, France; (19) The Ohio State University, Departments of Microbiology and Civil, Environmental and Geodetic Engineering, Columbus, Ohio, United States of America and The Ohio State University, Departments, Columbus, Ohio, United States of America; (20) Génomique Métabolique, Genoscope, Institut François Jacob, CEA, CNRS, Univ Evry, Université Paris-Saclay, Evry, France; (21) Oregon State University, Department of Microbiology, Oregon, United States of America; (22) Sorbonne Université, CNRS, Station Biologique de Roscoff, AD2M, UMR 7144, ECOMAP, Roscoff, France; (23) PANGAEA, Data Publisher for Earth and Environment Science, Bremen, Germany & MARUM—Center for Marine Environmental Sciences, Universität Bremen, Bremen, Germany; (24) Marine Biodiscovery Laboratory, School of Chemistry and Ryan Institute, National University of Ireland, Galway, Ireland; (25) Institut de Biologie de l’Ecole Normale Supérieure (IBENS), Ecole normale supérieure, CNRS, INSERM, Université PSL, Paris, France; (26) World Courier, an AmerisourceBergen Company, Russelsheim, Germany; (27) Division of Earth and Ocean Sciences, Duke University, Durham, North Carolina, United States of AMerica; (28) Laboratoire des Sciences de l’Environnement Marin (LEMAR), UMR 6539 UBO/CNRS/IRD/IFREMER, Institut Universitaire Européen de la Mer (IUEM), Brest, France; (29) Department of Earth Sciences, University of Southern California, Los Angeles, California, United States of America; (30) Marine Chemistry and Geochemistry Department, Woods Hole Oceanographic Institution; (31) Institut Pierre Simon Laplace, CNRS/IRD/MNHN (LOCEAN)Sorbonne-Université Paris Cedex 05, France; (32) Laboratoire Evolution, Génomes, Comportement et Ecologie, CNRS/Université Paris-Saclay, Avenue de la Terrasse, Gif sur Yvette, France. S. Planes and D. Allemand are joint scientific directors of Tara Pacific.

## Author Contributions

The study was conceived by all authors. The analysis approach was architected by Y.L. The specific methods and analysis were conducted by P.G. Data collection was led by E.B and G.B. Data processing was led by G.B. All authors interpreted the results, wrote the paper, and supported data analysis. TPC and TMM supported the extensive collection of the data and logistics.

