## Supplemental Material for "Emergent patterns of patchiness reflect decoupling between ocean physics and biology"

**Supplemental Methods**

All MODIS Aqua SST and chlorophyll-a was downloaded for August 2016. This period was selected both to overlap with the R/V Tara data and to conduct the analysis on data well before the known degradation of the MODIS-Aqua ocean color products (10.1117/12.2676873). SST and chl-a data were conservatively filtered using only data where both the chl-a and SST product flags indicated no warnings of bad data. The variance slope calculation was done on each individual image to preserve spatial patterns and these individual calculations of variance slope were then mean binned into 2° x 2° pixels.

**Supplemental Figures**

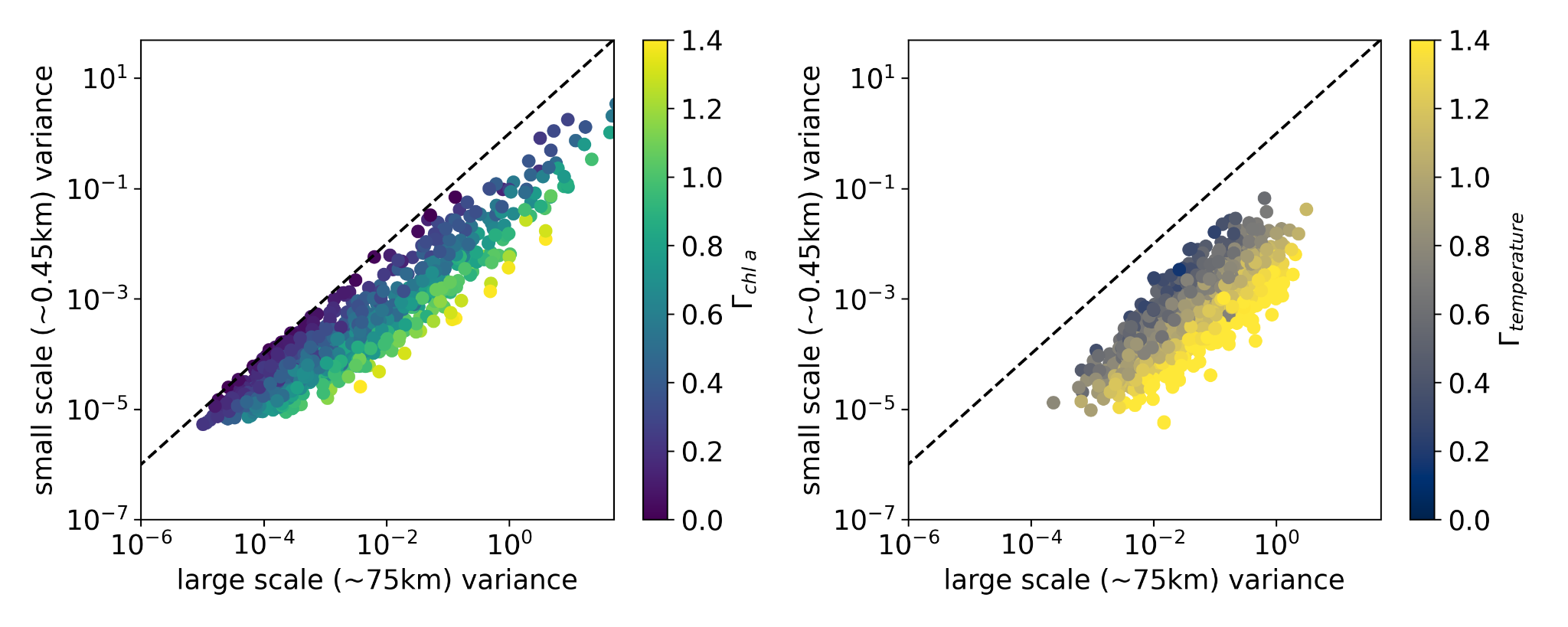

**Figure S1.** Large (75km) vs small scale (1km) variance colored by the variance slope (𝛤) for chlorophyll-a and temperature (left).

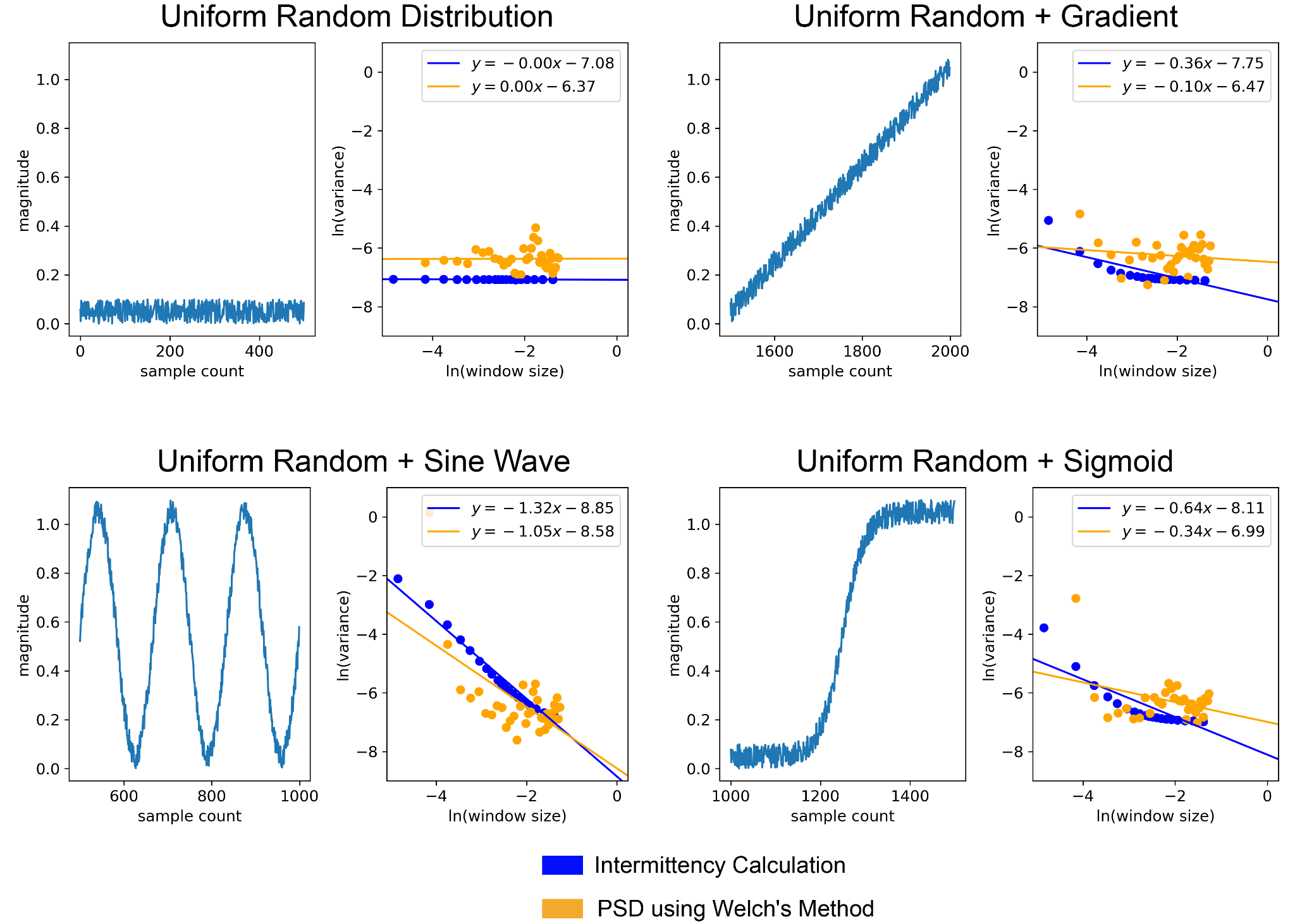

**Figure S2.** Comparisons of variance slope (𝛤) and a typical power spectral density slope calculated via Welch’s Method for four different synthetic patterns.

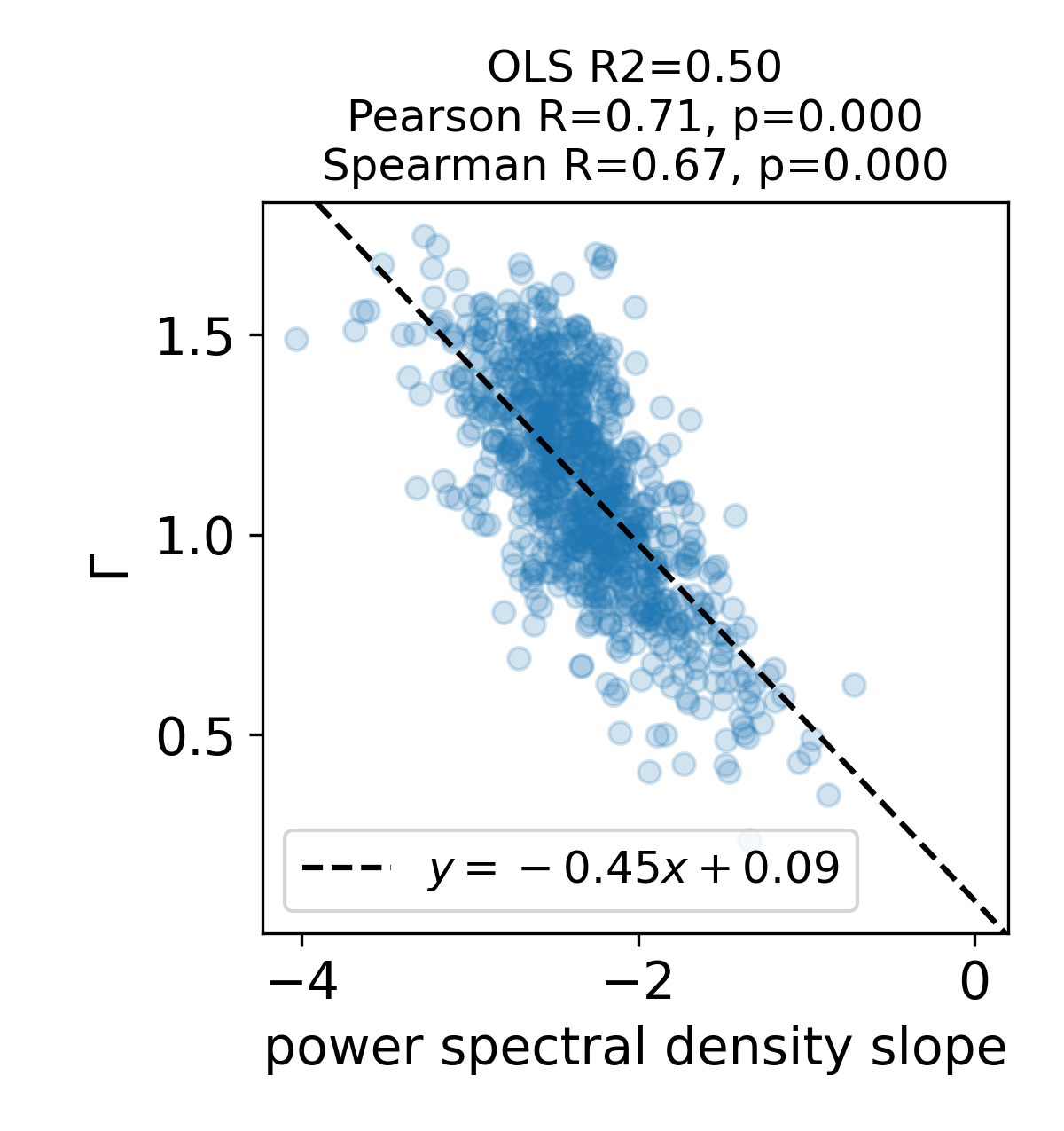

**Figure S3.** The relationship of variance slope (𝛤) and the power spectral density slope for a large dataset of temperature collected on the R/V *Tara* showing the tight correspondence and the formula for conversion between the two.

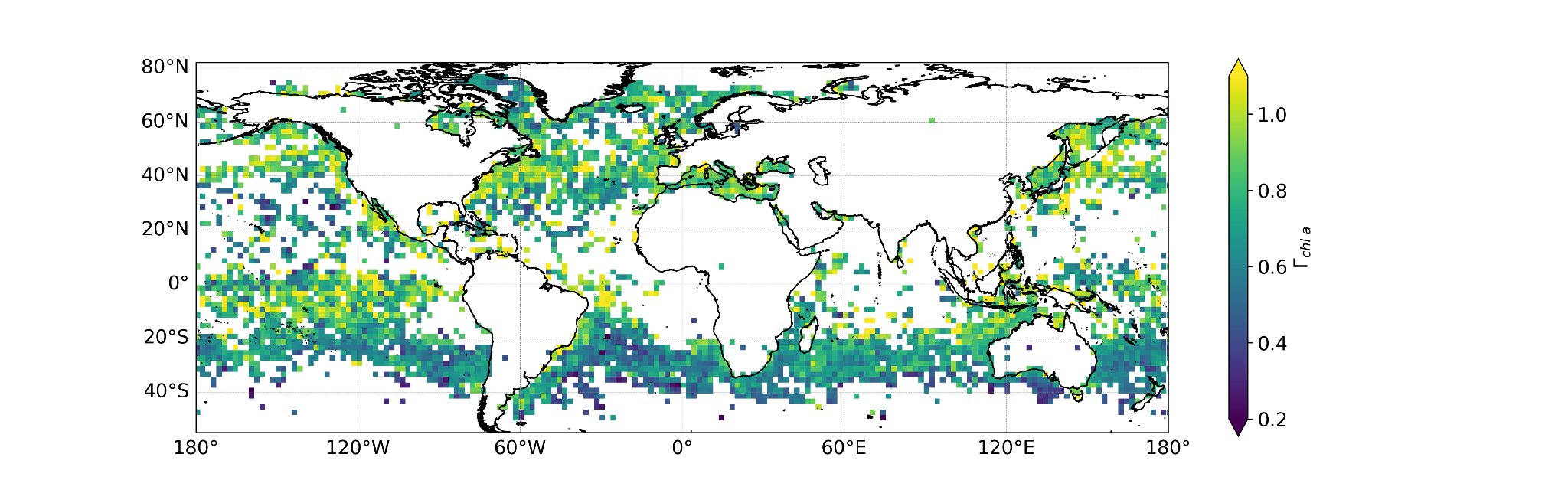

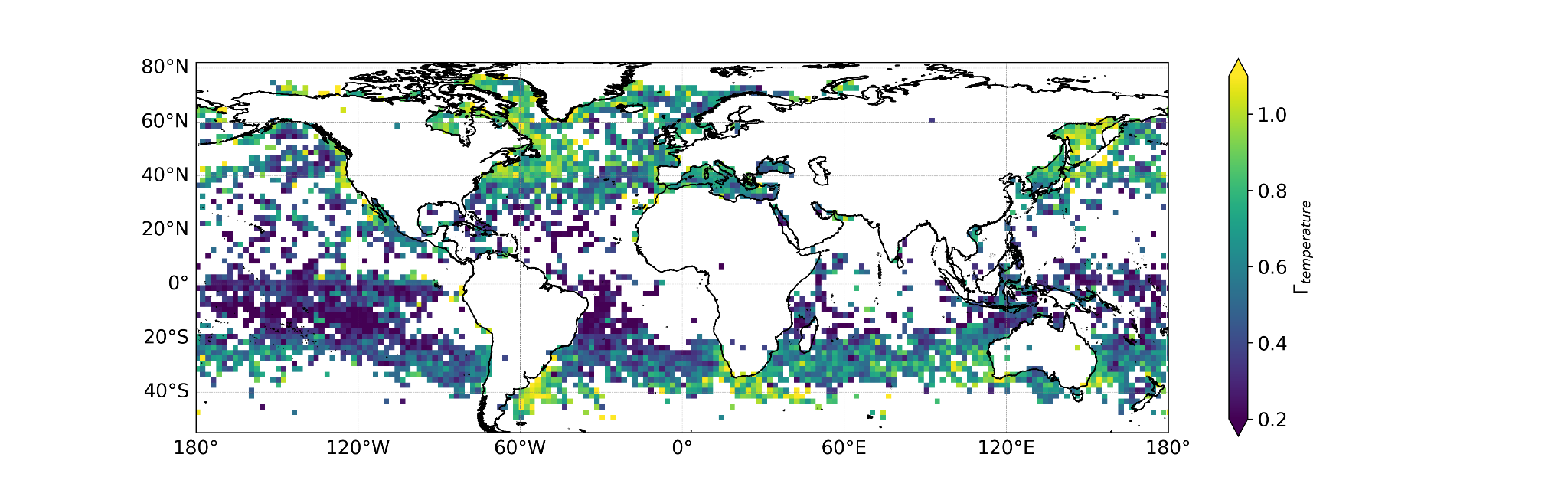

**Figure S4.** Variance slope (𝛤) for chlorophyll-a (top panel) and temperature (bottom panel). Note that the calculation here is done from 3 km to 75 km (dx=1) and then interpolated to a 2x2° grid where all slope values within each grid square are averaged. This was to match as closely as possible the R/V *Tara* analysis which goes from .45 km to 75 k (dx=0.15km)

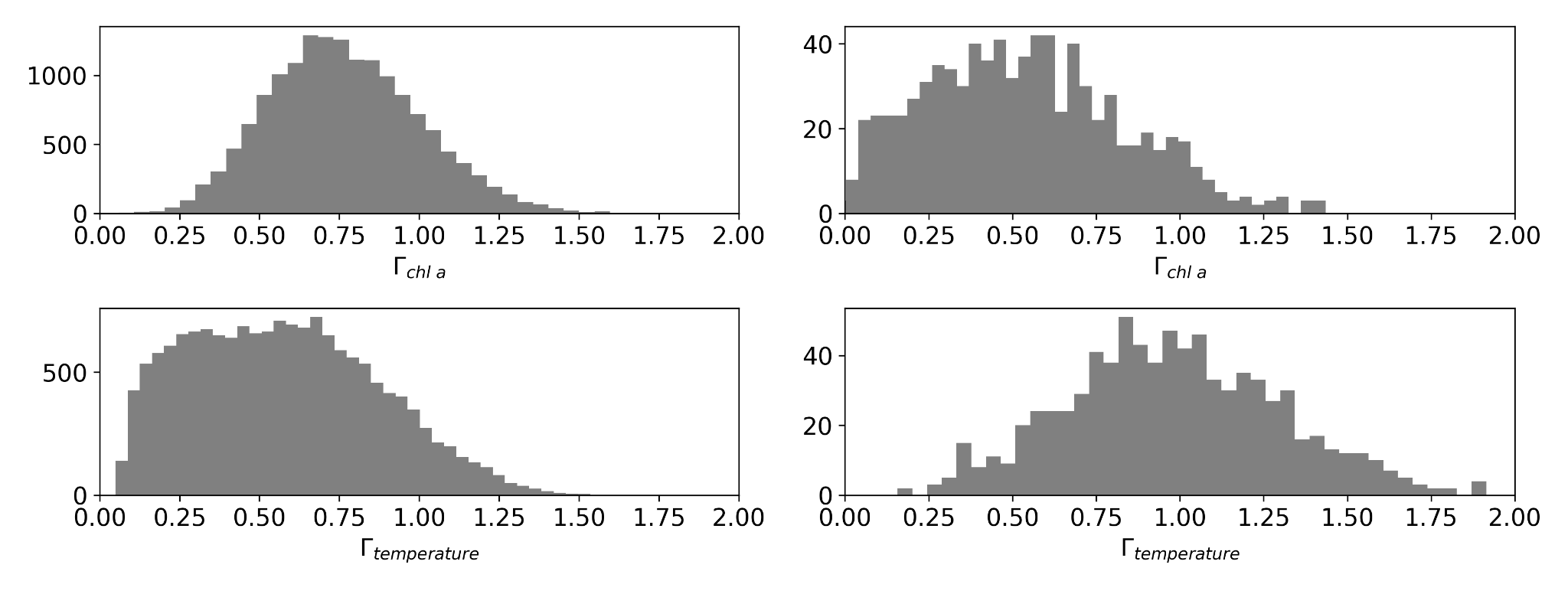

**Figure S5.** Satellite (left) and R/V *Tara* (right) chlorophyll-a and temperature variance slopes (𝛤). The satellite data is from MODIS Aqua and consists of data from all of Aug 2016.

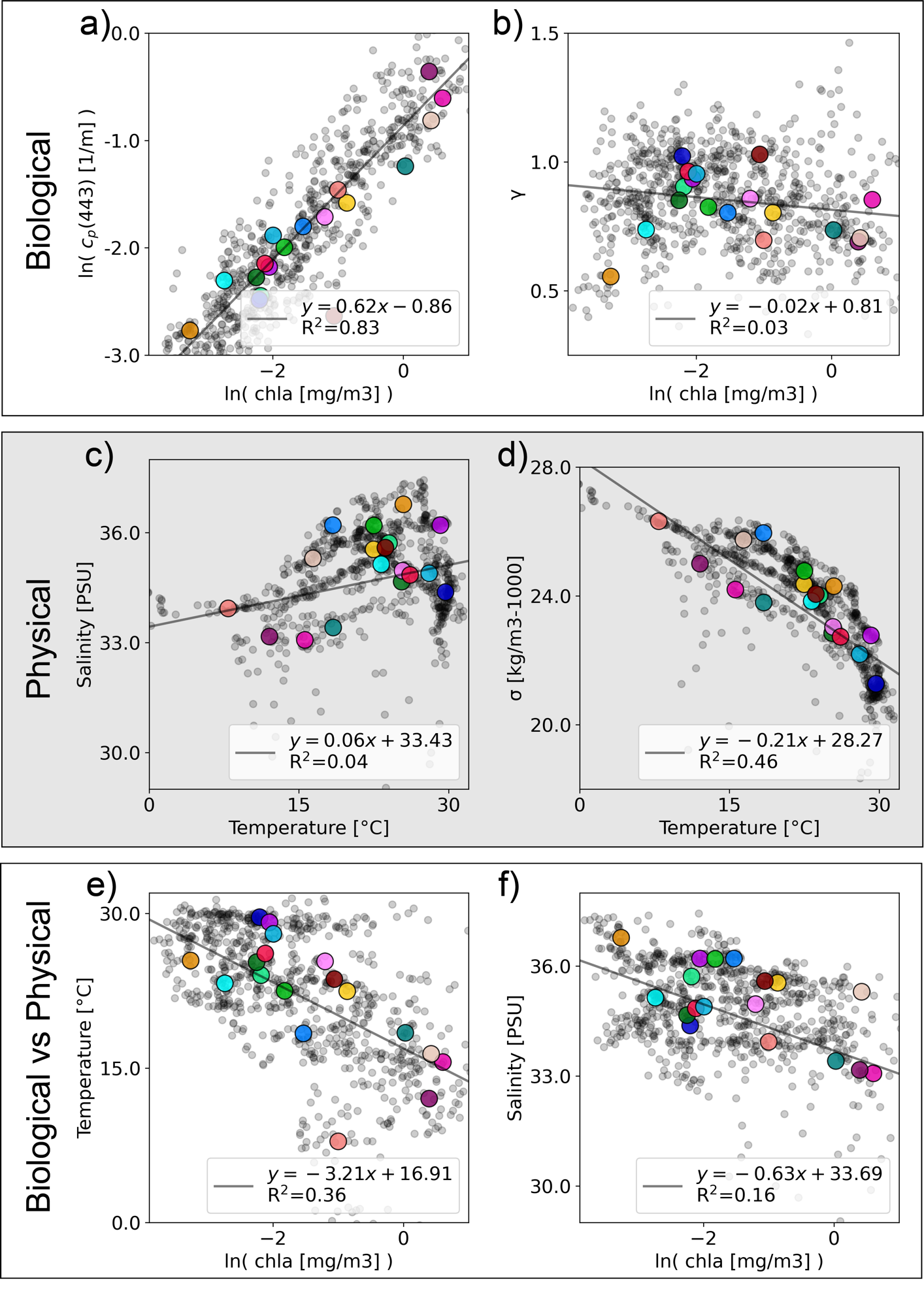

**Figure S6.** Absolute value comparisons between variables across the dataset. Larger colored markers correspond with the Longhurst provinces from Figure 5. N.b. correlations are shown for all data, not the Longhurst province means, and all plots have a p-value < 0.001.

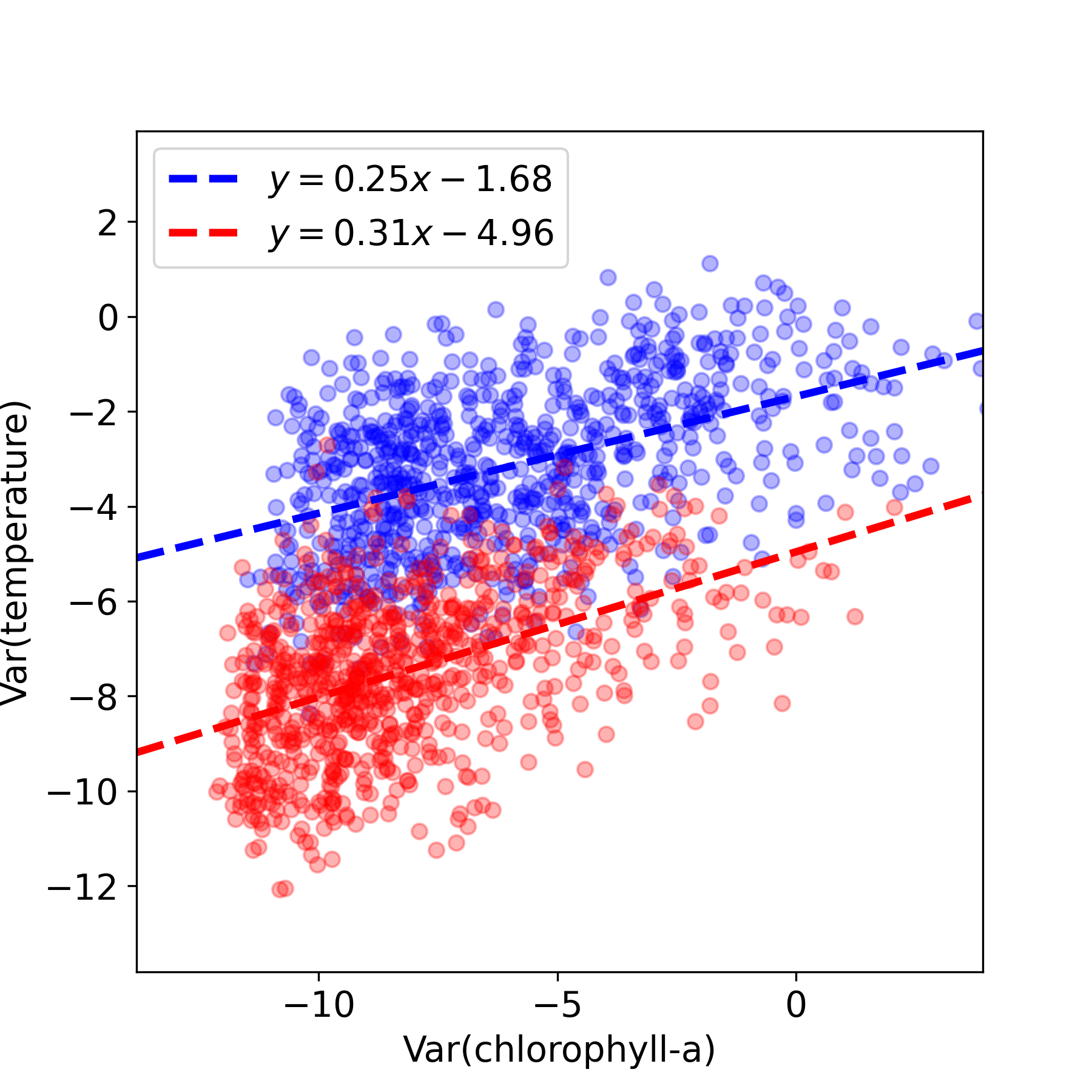

**Figure S7.** Temperature variance as a function of chlorophyll variance at the large scale (~75km, blue) and at the small scale ( ~0.45km, red). While the variances do have a correlation (R^2^=0.22 in both cases), the variance slope (𝛤) does not correlate between physical and biological variables (R^2^0.03).
